# In-scanner head motion and structural covariance networks

**DOI:** 10.1101/2021.04.28.441807

**Authors:** Heath R. Pardoe, Samantha P. Martin

**Author notes:** Corresponding Author: Heath Pardoe, PhD, Department of Neurology, NYU Grossman School of Medicine, 145 East 32nd St, 5^th^ Floor, room 507, New York City, NY 10016, USA.

## Abstract

In-scanner head motion systematically reduces estimated regional gray matter volumes obtained from structural brain MRI. Here, we investigate how head motion affects structural covariance networks that are derived from regional gray matter volumetric estimates. We acquired motion-affected and motion-free whole brain T1-weighted MRI in 29 healthy adult subjects and estimated relative regional gray matter volumes using a voxel-based morphometry approach. Structural covariance network analyses were undertaken while systematically increasing the number of motion-affected scans included in the analysis. We demonstrate that the standard deviation in regional gray matter estimates increases as the number of motion-affected scans increases. This subsequently increases pair-wise correlations between regions, a key determinant for construction of structural covariance networks. We further demonstrate that head motion systematically alters graph theoretic metrics derived from these networks. Our findings suggest that in-scanner head motion is a source of error that violates the assumption that structural covariance networks reflect neuroanatomical connectivity between brain regions. Results of structural covariance studies should be interpreted with caution, particularly when subject groups are likely to move their heads in the scanner.

## 1. Introduction

Analysis of the covariance of regional grey matter volume and related morphometric measures across subjects is a widely used MRI-based network analysis technique that is used to investigate neuroanatomical changes in disease and healthy aging [1–15]. The primary underlying assumption of structural covariance analyses is that regional covariation of morphometric estimates reflects underlying neuroanatomical connectivity between brain regions. In this study we test this assumption by investigating the effect of in-scanner head motion on construction and graph theoretic analysis of structural covariance networks.

In-scanner head motion has been previously demonstrated to cause brain-wide changes in morphometric estimates derived from structural MRI scans [16–19]. Increased head motion is generally associated with brain-wide reductions in cortical thickness and regional gray matter volume. Previous studies have shown that head motion is typically increased in disease groups relative to healthy controls and varies by age, with children and elderly adults showing increased head motion relative to adults aged between 15 and 40 years [17, 19]. Although the effect of head motion on MRI-based estimates of regional cortical thickness, volume and related neuroanatomical measures has been extensively described recently, the effect of motion on higher order multivariate analytical approaches such as structural covariance analysis is yet to be established.

Structural covariance networks are inferred by measuring the correlation of brain volumes between pairs of brain regions across subjects. Because head motion increases variability in regional morphometric estimates across the whole brain, it is likely that the variability in both members of any pair of regional estimates is increased, increasing the measured correlation between these pairs. This effect can be conceptualized as a corollary of the restricted range phenomenon, in which the measured correlation between two variables is reduced if the sample is drawn from a limited range of the population [20]. In the case of in-scanner head motion, we hypothesize that the inclusion of motion affected scans will increase variability in morphometric estimates across subjects and subsequently increase the measured correlation in volumes between pairs of brain regions.

We investigated the effect of head motion on structural covariance networks by obtaining both motion-affected and motion-free whole brain T1-weighted MRI scans in healthy adult subjects. MRI scans were used to obtain regional gray matter volume estimates following a voxel-based morphometry approach similar to that used in a number of prior studies [4, 7, 8, 11, 12, 15]. We systematically varied the number of motion affected scans included in each group and assessed how this influenced metrics related to network construction and commonly used graph theoretic measures. If the inclusion of motion-affected scans affects the construction of structural covariance networks and derived metrics, regional covariance of volumes and related morphometric measures across subjects may not always reflect anatomical connectivity. In general, if subject groups differ in the amount of in-scanner head motion, differences in graph theoretic network metrics derived from structural covariance analyses of these groups may not be due to neurobiological differences.

## 2. Methods

Whole brain T1w MRI scans were obtained in 29 healthy control subjects recruited via community advertisement (mean age 33 ± 13 years, 13 female). Approval for the study was obtained from the NYU Langone Health Institutional Review Board, and written informed consent was obtained from all study participants. MRI scans and scripts associated with the analyses presented in this study are provided at https://sites.google.com/site/hpardoe/structcovar_motion [21].

Participants were imaged on a Siemens 3T Prisma MRI scanner. Two T1w MPRAGE MRI scans were acquired per participant in each imaging session; a motion affected scan, in which participants moved their heads in stereotyped prompted motions during the scan, and a motion-free scan in which participants held their head still. Image acquisition parameters were: 1mm isotropic voxel size, echo time (TE) = 3 ms, flip angle = 8 degrees, inversion time (TI) = 0.9 s, repetition time = 2.5 s.

MRI scans were segmented and co-registered using the DARTEL toolbox provided in the SPM12 software package [22]. A custom template was created from the motion-free dataset. Scans were segmented into gray (GM) and white matter and coregistered to MNI space via non-rigid coregistration to the custom template. GM segments were smoothed using an 8mm Gaussian smoothing filter. Regional GM volumes were estimated by calculating the mean GM value within each region defined in the AAL atlas [23]. We excluded cerebellar brain regions from our analyses, thereby yielding GM volume estimates in 90 brain regions. Covariance matrices were estimated from regional GM values across subjects by measuring the Pearson correlation (r) between brain regions. Matrices were thresholded and binarized to construct undirected adjacency matrices. Undirected adjacency matrices were thresholded by calculating the Pearson r value threshold that corresponded to a fixed target edge density; this is a standard approach used in a number of previous studies [2, 3, 6, 9, 13]. Subject groups were formed by systematically increasing the number of motion-affected scans in each group, ranging from no motion affected scans to all motion affected scans.

We investigated how in-scanner head motion affected structural covariance network construction by measuring the following as the number of motion-affected scans included in each dataset increased: (i) the variability in volume estimates, assessed by measuring the standard deviation in each brain region across subjects; (ii) Pearson correlation coefficients (r) between volumes of brain regions and (ii) the Pearson r thresholds required to obtain fixed edge densities = 0.1,0.15 and 0.2. We also investigated how commonly used graph theoretic network metrics, consisting of (i) global clustering coefficient (transitivity), (ii) modularity, (iii) global efficiency and (iv) small world index varied as a function of the number of motion-affected scans in each group. These analyses are described in more detail below. Analyses were carried out using the R software package with the iGraph, brainGraph and qgraph libraries [24–27].

A bootstrapping approach was used to estimate how GM estimates derived from motion-affected scans affected (i) the across-subject standard deviation in each brain region and (ii) the Pearson correlation r between pairs of brain regions. The number of motion affected scans included was systematically increased from zero (no motion affected scans) to 29 (all motion affected scans). For each iteration, N = 5000 samples were drawn with replacement and (i) the across-subject standard deviation in each brain region and (ii) pairwise correlations in mean GM values between brain regions were calculated. The relationship between average standard deviation across brain regions and the percentage of motion affected scans was tested using a linear model. The relationship between average pairwise correlation and the percentage of motion affected scans was tested using a linear model.

We further investigated how the inclusion of motion-affected scans affected the threshold r value required to obtain undirected adjacency matrices with a pre-specified fixed edge density. Target edge densities of 0.1, 0.15 and 0.2 were used. As in previous analyses a bootstrapping approach was used to estimate the average r value required to obtain the target densities as the number of motion-affected scans increased. 5000 samples were drawn with replacement for each iteration as the number of motion-affected scans in the dataset was increased. The relationship between threshold r value and percentage of motion affected scans was tested using a linear model.

Finally we investigated how in-scanner head motion affects graph theoretic network measures that are commonly used in structural covariance studies to assess differences between subject groups. Specific metrics included measures of network segregation, consisting of transitivity (also known as clustering coefficient) and modularity, and measures of network integration including global efficiency. We also measured how motion-affected scans influence the measured small world index. For these analyses the graph theoretic estimates were obtained from graphs constructed from adjacency matrices that were thresholded to obtain target edge densities = 0.1, 0.15 and 0.2. As in the previous analyses 5000 samples were drawn with replacement as the number of motion affected scans increased, with the four graph theoretic metrics calculated for each sample.

## 3. Results

The average across-subject standard deviation systematically increased as the number of motion-affected scans included in an analysis increased, with an increase of 1.9 × 10^−4^ SD per percent change in the number of motion-affected scans (p = 4.5 × 10^−10^). The change in standard deviation as a function of the number of motion affected scans, both per-region and averaged across all brain regions, is shown in Figure 1. For example, if 20% of the scans in a dataset were motion affected, the standard deviation increased on average by 23%. The average correlation between GM regions also systematically increased as the number of motion-affected scans included increased, with an increase of 5.1 × 10^−4^ in r per percent change in the number of motion-affected scans (p = 1.5 × 10^−6^, Figure 2). As an example, we have provided scatterplots showing the change in the correlation between the left pallidum and left inferior temporal gyrus as the number of motion affected scans increased (Figure 3). The figure shows that the Pearson correlation increased from r = 0.57 if no motion affected scans are included to r = 0.71 if 6 subjects (20% of the sample) are motion affected. The average threshold r value required to obtain adjacency matrices with the pre-specified edge densities systematically increased as the number of motion affected scans included in the analyses increased; this was observed across the three target densities investigated in our study (Figure 4).

**Figure 1.**
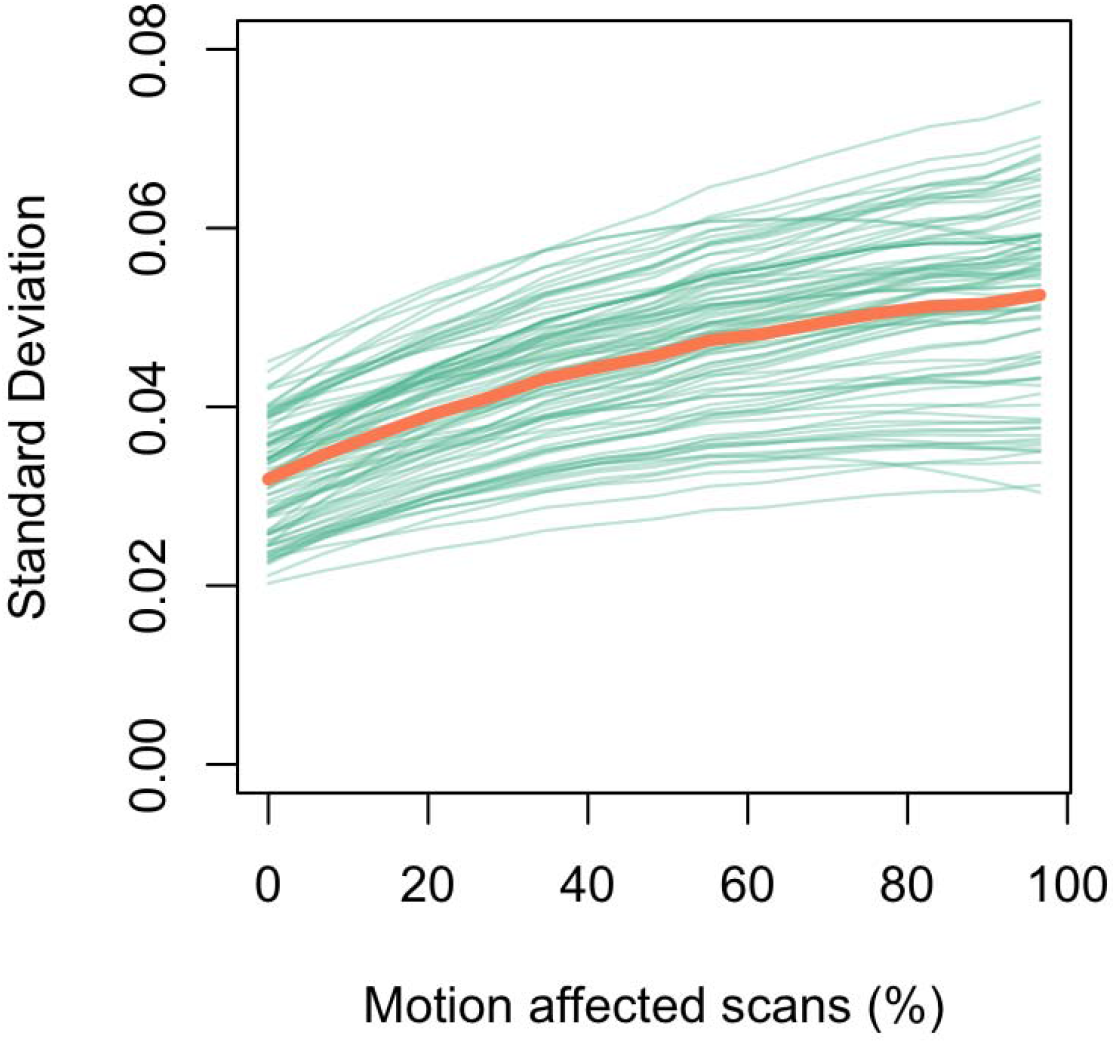
Variability in regional gray matter volume estimates across subjects increases with the number of motion affected scans included in a dataset. The standard deviation averaged across all brain regions is shown in orange, with standard deviation in individual brain regions shown in green.

**Figure 2.**
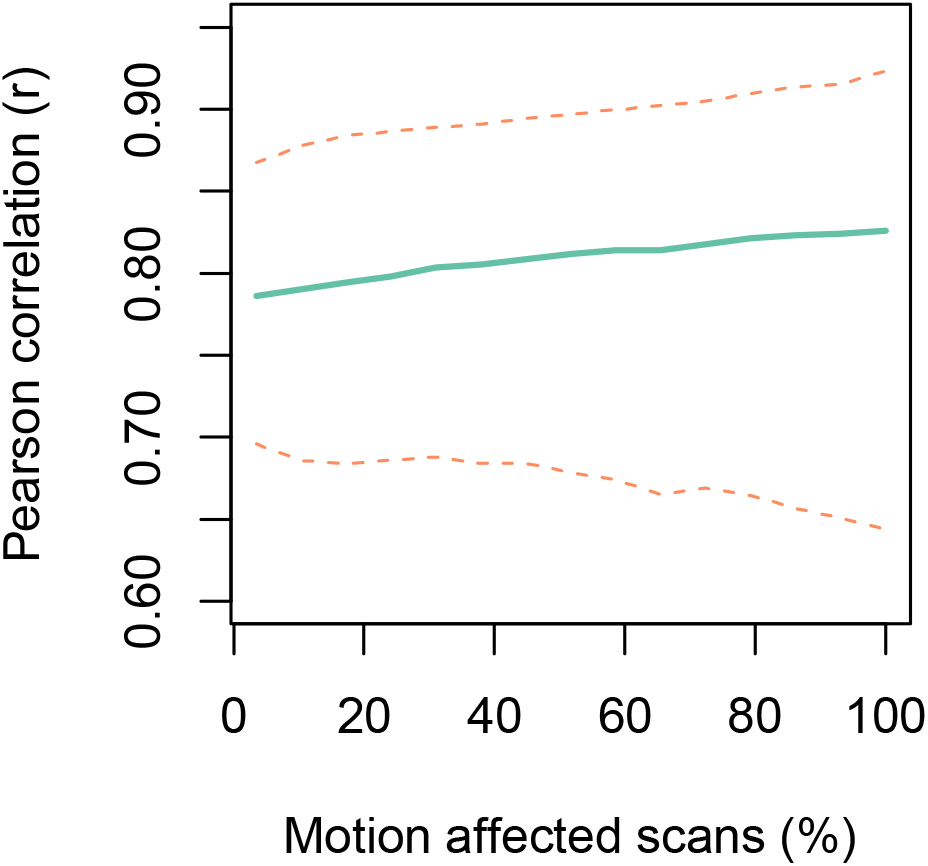
The correlation between gray matter volume estimates increases as the number of motion-affected scans included in a dataset increases. The green line shows the average Pearson r across all brain regions and the dashed lines indicate the 95% confidence interval.

**Figure 3.**
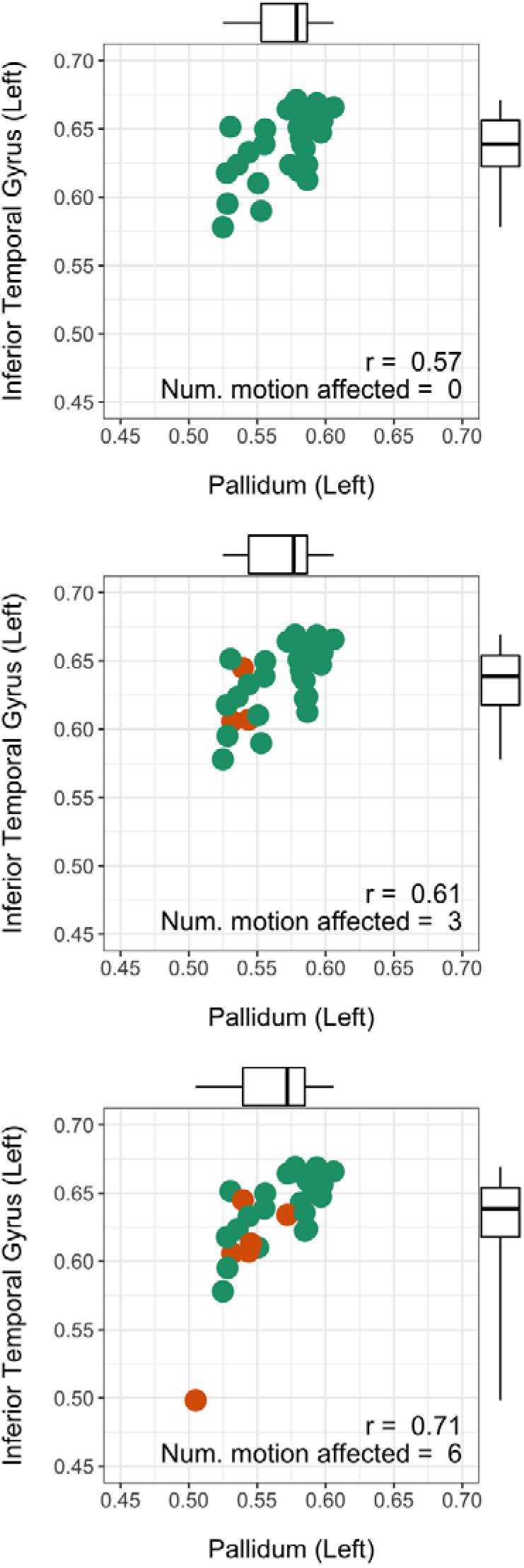
The effect of in-scanner head motion on the measured correlation in gray matter volume between the pallidum and the inferior temporal gyrus. As more motion-affected scans are included in the sample the correlation between these brain regions increases from 0.57 to 0.71.

**Figure 4.**
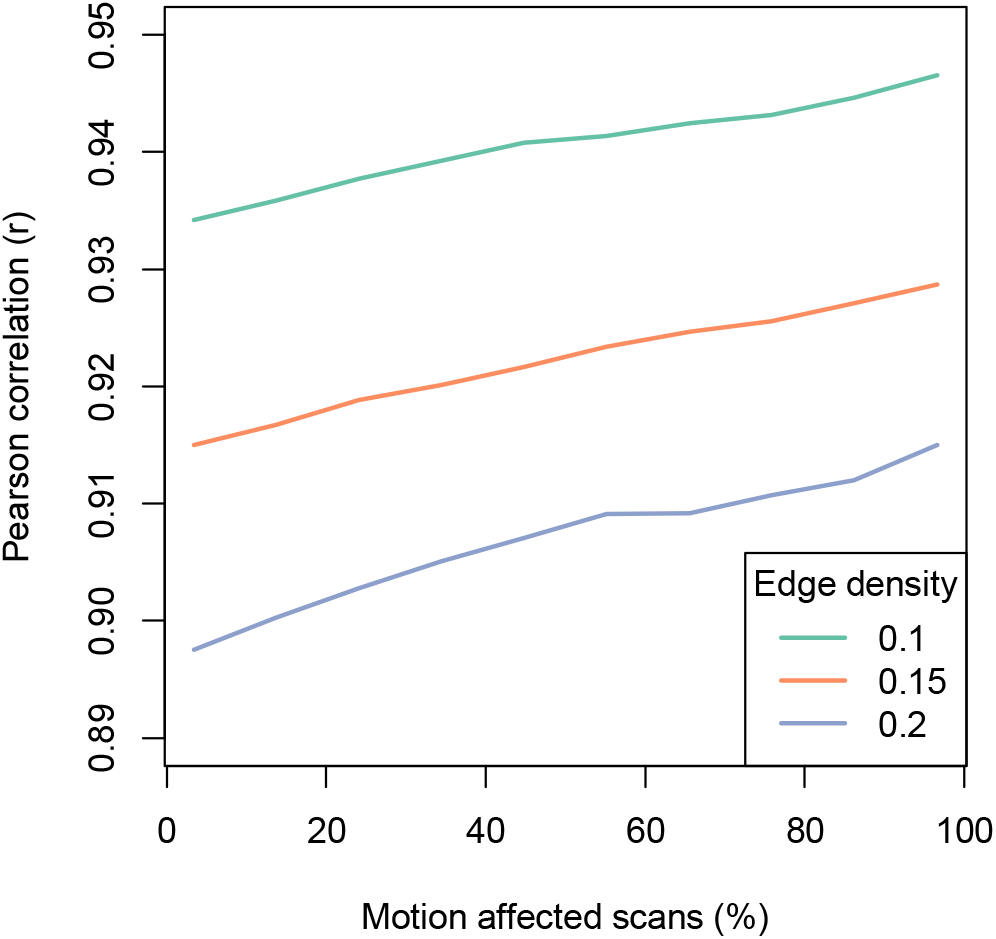
The threshold Pearson correlation required to obtain a pre-defined network edge density is dependent on the number of motion-affected scans included in an analysis.

Both modularity and efficiency systematically increased as the number of motion affected scans increased. These changes were consistent across the range of edge densities used in our analyses (Figure 5). Small world index generally decreased as the number of motion affected scans increased, although the higher edge densities = 0.15, 0.2 showed a small peak in small world index when a low number of motion affected scans was increased. The relationship between transitivity and number of motion affected scans depended on edge density, with the shape of the density = 0.1 curve markedly different to curves for edge densities = 0.15 and 0.2. For the higher edge density values the transitivity decreased as the number of motion affected scans increased. When examining changes in graph theoretic metrics for a single edge density = 0.15, it can be seen that the most substantial changes in transitivity, modularity and efficiency occur when a small number of motion affected scans are included in the analysis (Figure 6).

**Figure 5.**
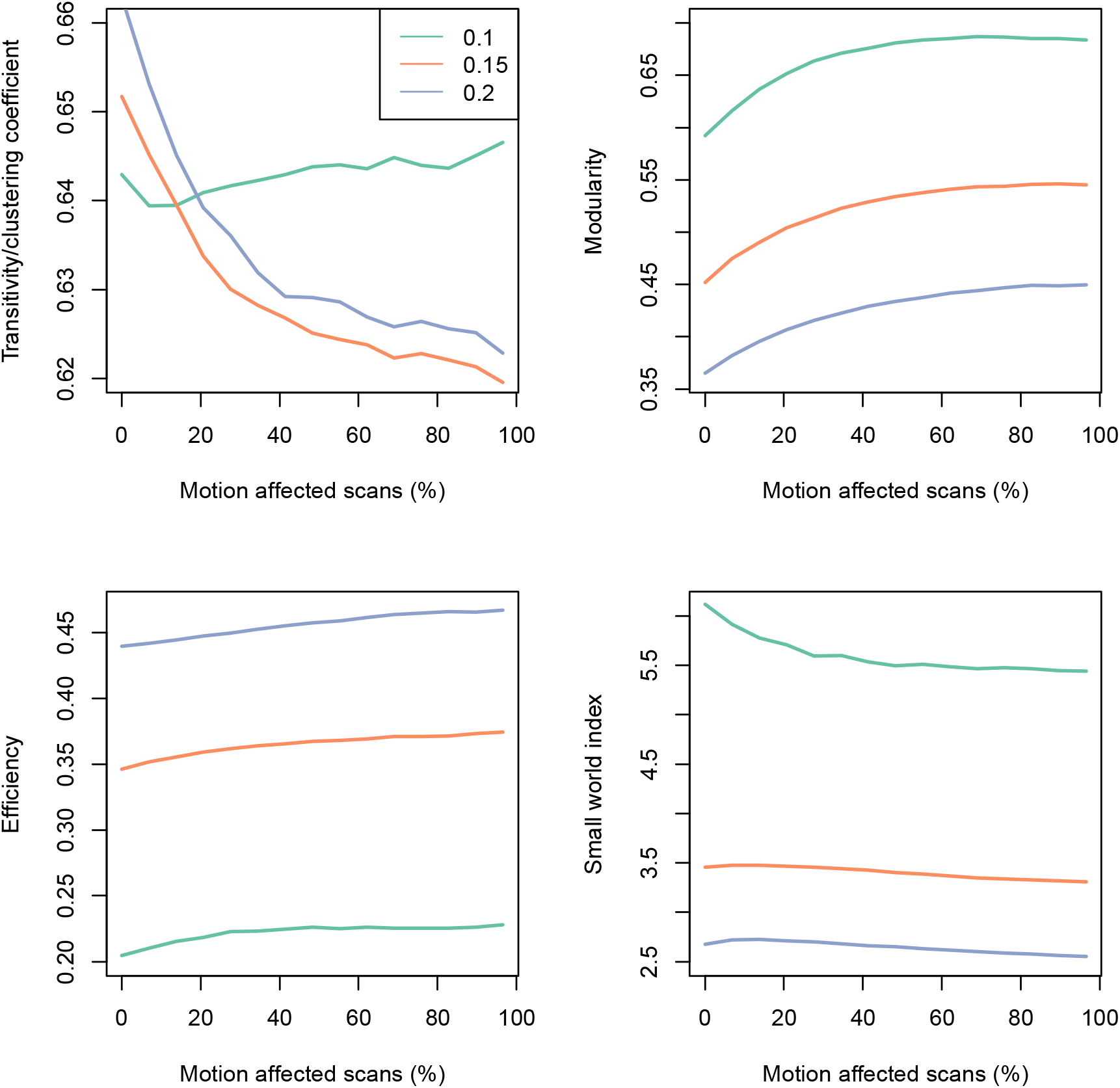
Change in graph theoretic metrics across edge density = 0.1, 0.15, 0.2. The relationship between motion and transitivity (clustering coefficient) is variable depending on the edge density used. Modularity and efficiency increase as the number of motion affected scans increases. Small world index generally decreases as the number of motion affected scans increases.

**Figure 6.**
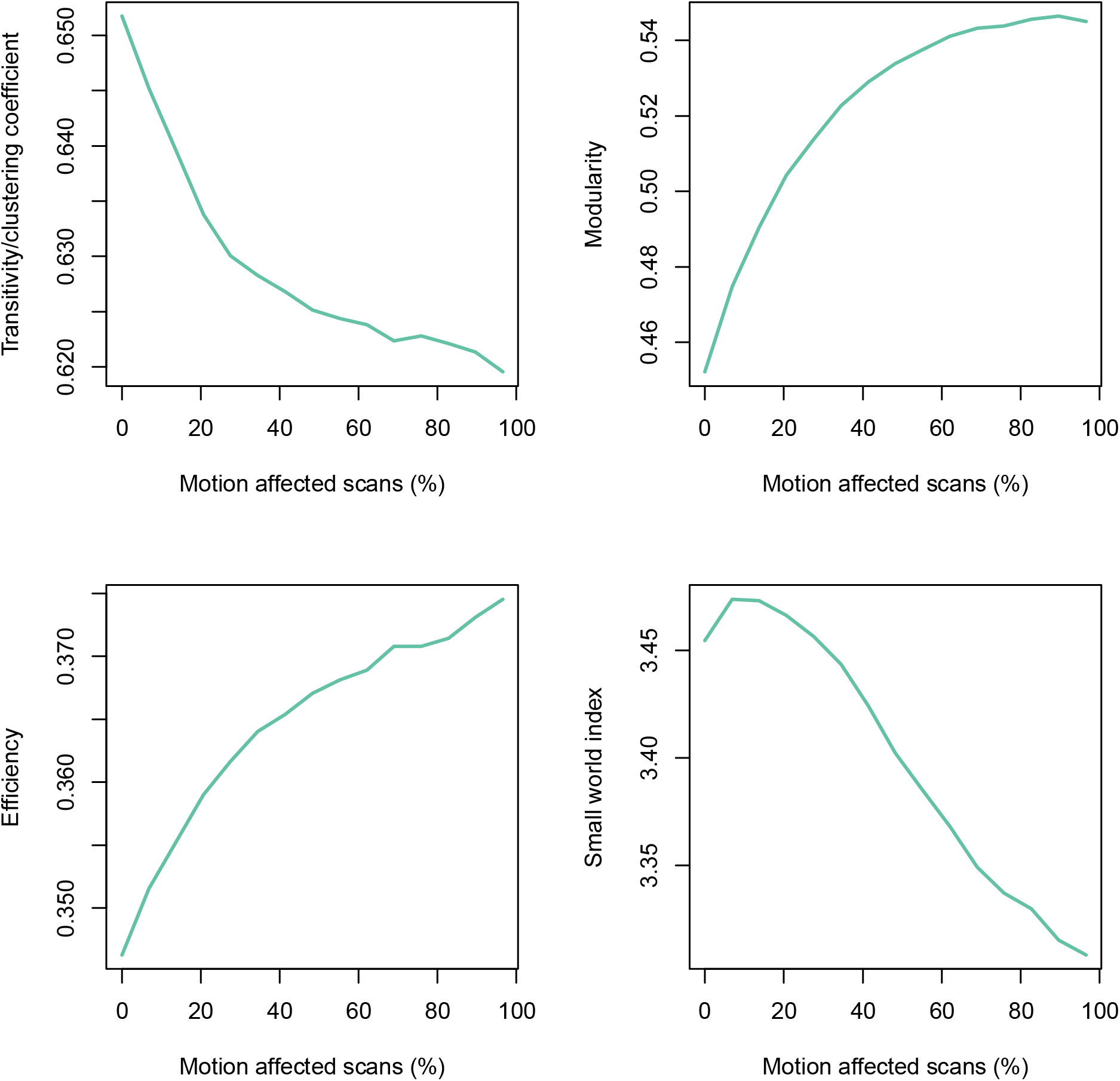
Change in graph theoretic metrics as a function of in-scanner motion for edge density = 0.15. For transitivity, modularity and efficiency the largest changes occurred when a small number of motion-affected scans were included (< 20%).

## 4. Discussion

We have demonstrated that the inclusion of motion-affected scans in a structural covariance analysis framework causes systematic increases in the variability of measured gray matter volumes. The increase in variability subsequently increases the correlation in GM volumes between brain regions. The increases in regional variability and measured correlation between brain regions are observed across the whole brain. These findings suggest that in-scanner head motion can affect the construction of networks derived from covariance of the volumes of gray matter regions across subjects. We have also shown that graph theoretic metrics derived from graphs constructed based on correlation between brain regions are dependent on the number of motion affected scans included in an analysis. Our analyses suggest that in-scanner head motion is a potential source of error for structural covariance analyses, and violates the primary underlying assumption that significant correlations in GM volume estimates between brain regions reflect neuroanatomical connectivity. These findings suggest that caution should be exercised when interpreting the results of structural covariance analyses.

Our experimental framework used individuals who deliberately moved their heads in the MRI scanner. The motion-related artifacts in these scans may be greater than those encountered in a typical clinical imaging study, particularly in studies that have stringent quality control procedures. Although quality control protocols are likely to reduce the magnitude of this source of error, it is unclear if they would eliminate this effect. Previous studies have demonstrated that in-scanner head motion in a clinical setting has a significant effect on cortical thickness and regional volume even in scans that have been visually rated as being high quality [17]; this suggests that it is possible that even subtle in-scanner head motion could have a systematic effect on structural covariance analyses, particularly in large studies. Recently developed motion-robust acquisitions [28] are likely to ameliorate the effects of in-scanner head motion described in this work, however this should be explicitly investigated in future studies.

In summary, we have demonstrated that in-scanner head motion systematically increases the correlation in estimates of regional gray matter volume as a consequence of increased variability in these estimates. These changes affect graph theory-based metrics typically used to infer differences in neuroanatomical connectivity between subject groups. As with mass-univariate analyses of regional brain volume differences between subject groups, the results of structural covariance analyses should be interpreted with the knowledge that head motion is a potential source of error.

## Acknowledgements

This work was supported by FACES Foundation, NY USA

